# GenePert: Leveraging GenePT Embeddings for Gene Perturbation Prediction

**DOI:** 10.1101/2024.10.27.620513

**Authors:** Yiqun Chen, James Zou

**Affiliations:** Department of Biomedical Data Science, Stanford University, Stanford; Departments of Biomedical Data Science, EE, and CS, Stanford University, Stanford

**Keywords:** single-cell RNA Sequencing, Gene Perturbation, AI for Biology, regularized regression

## Abstract

Predicting how perturbation of a target gene affects the expression of other genes is a critical component of understanding cell biology. This is a challenging prediction problem as the model must capture complex gene-gene relationships and the output is high-dimensional and sparse. To address this challenge, we present GenePert, a simple approach that leverages GenePT embeddings, which are derived using ChatGPT from text descriptions of individual genes, to predict gene expression changes due to perturbations via regularized regression models. Benchmarked on eight CRISPR perturbation screen datasets across multiple cell types and five different pretrained gene embedding models, GenePert consistently outperforms all the state-of-the-art prediction models measured in both Pearson correlation and mean squared error metrics. Even with limited training data, our model generalizes effectively, offering a scalable solution for predicting perturbation outcomes. These findings underscore the power of informative gene embeddings in predicting the outcomes of unseen genetic perturbation experiments *in silico*. GenePert is available at https://github.com/zou-group/GenePert.

## 1 Introduction

Understanding the function, organization, and underlying mechanisms of cellular systems is critical to advancing biology and rationally manipulating cells. This has remained challenging for decades because most high-throughput data have been observational in nature, meaning we could not establish causality. A series of recent breakthroughs in experimental and computational biology has brought us one step closer to understanding these issues. On the experimental side, Perturb-seq combines pooled genetic perturbation screens in cells and tissues (such as CRISPR-Cas9, CRISPRi, and CRISPRa, which knock out, inhibit, and activate transcription, respectively) with high-content single-cell readouts, such as RNA or chromatin profiles, providing *interventional* data [Dixit et al., 2016, Rood et al., 2024, Bock et al., 2022]. However, given the large number of genes in the genome, as well as the number of possible gene perturbations, large-scale Perturb-seq screens remain expensive [Yao et al., 2023]. This has spurred much interest and advances on the computational side, where machine learning models are trained on data from these high-throughput perturbation screens to predict responses to unseen genetic perturbations [Roohani et al., 2024, Lotfollahi et al., 2020]. This presents a challenging prediction problem, as the output can be high-dimensional (ranging from 1,000 to 20,000) and sparse, while the number of unique perturbations is comparatively smaller (ranging from 20 to 1,500).

Pioneering work in *in silico* perturbation prediction used linear models on the perturbation feature matrix (i.e., a matrix with 1 for the perturbed gene and 0 otherwise) to describe the mean impact of each perturbation on gene expression across cells [Dixit et al., 2016, Duan et al., 2019, Yang et al., 2020]. However, the linearity in the raw perturbation feature space makes it difficult to predict unseen perturbations, and performance is limited by the linear assumption [Lotfollahi et al., 2020]. This has inspired the adoption of deep learning models to generate non-linear representations of the input data that are more robust to non-perturbation-related variations, such as technical and biological noise in experiments. For example, scGen [Lotfollahi et al., 2019] learned a low-dimensional latent space using a variational autoencoder, modeling how perturbations impact this latent space. The trained model can extrapolate the effect of new perturbations on test cells by first computing the latent space and then decoding it back into gene expression space. CPA [Lotfollahi et al., 2020], a follow-up work, refines this idea by incorporating interpretable linear models within the latent embedding space. As gene-gene interactions are often of central interest in perturbation screens, GEARS, often considered state-of-the-art, employs graph neural networks to encode prior knowledge of gene-gene relationships, enabling the prediction of the effects of perturbing multiple genes unseen during training. More recently, with the rise of large-scale pretrained “foundation models” for single-cell transcriptomics [Cui et al., 2024, Theodoris et al., 2023, Hao et al., 2024], studies have shown state-of-the-art predictive performance by fine-tuning these learned embeddings for perturbation screen experiments. The intuition is that these pretrained gene and cell embeddings offer better contextual representations for perturbation screen data than previous methods, which often had to learn latent representations from scratch. Despite the success of deep learning-based models, recent studies have revealed that very simple baselines, such as averaging the training examples for unseen perturbations, can outperform these complex models on benchmark datasets [Märtens et al., 2024, Kernfeld et al., 2024, Ahlmann-Eltze et al., 2024, Wenteler et al., 2024, Gaudelet et al., 2024]. A compelling hypothesis is that high-dimensional and sparse single-cell Perturb-seq data do not always have the signal-to-noise ratio necessary to support intricate non-linear relationship modeling. Therefore, simpler models, while theoretically more biased due to over-simplification of the relationships, have less variance in their predictions. Motivated by the observation of the bias-variance trade-off, in this work, we propose GenePert, a simple 𝓁_2_-regularized linear regression model that captures gene perturbations using pretrained gene embeddings (see Figure 1).

**Figure 1:**
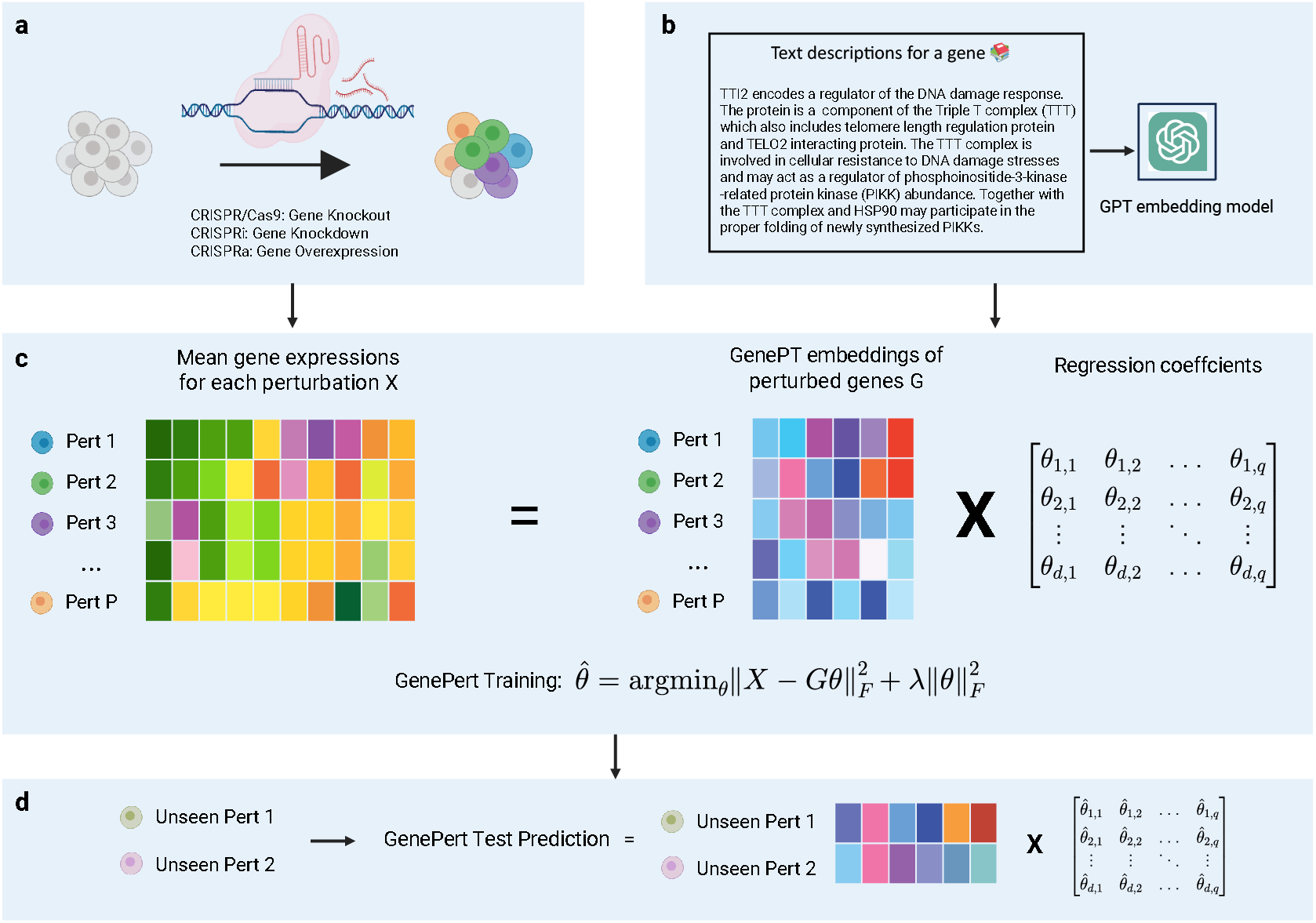
Overview of GenePert for perturbation screen data prediction. (a) Data from pooled perturbation screens with high-throughput readouts (for instance, single-cell RNA sequencing) serve as input for the GenePert model. (b) For each gene being perturbed, we use its GenePT embedding as its feature: that is, we first extract its corresponding gene information summary from NCBI and, if available, its protein summary from UniProt, and use OpenAI’s Model 3 text embedding of the summary as its representation. (c) For training, GenePert uses ridge regression, which posits a linear relationship between the average regulatory effects (left matrix *X*) of perturbations (rows) and the gene features (right matrix *G*). (d) During testing, for an unseen perturbation, GenePert uses the fitted coefficients 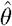 on the corresponding GenePT embedding to generate predictions.

More specifically, we represent perturbed genes using their GenePT gene embeddings [Chen and Zou, 2023], which are large language model embeddings of descriptions of individual gene facts and functionalities. These embeddings have been demonstrated to be effective encoders of biological knowledge, particularly in the context of gene-gene interactions and functionality predictions. We also propose a simple method to integrate training and prediction for double gene perturbations by summing their corresponding GenePT embeddings, which leads to predictive performance that outperforms other methods.

Compared to prior and concurrent work employing similar simple, off-the-shelf predictive models on top of gene embedding strategies [Märtens et al., 2024, Gaudelet et al., 2024, Ahlmann-Eltze et al., 2024, Wenteler et al., 2024], our approach achieves better prediction results and more naturally generalizes to unseen gene perturbations. This is because GenePT embeddings can be easily generated from textual descriptions of genes, rather than requiring (i) observing the gene expression profile to extract principal components, or (ii) extensive pre-training like other single-cell foundation models. Additionally, the linear combination of single-gene perturbation effects to predict double-gene perturbation effects has been concurrently explored by Gaudelet et al. [2024] and Ahlmann-Eltze et al. [2024]. In contrast to their work, our approach does not rely on observing at least one perturbation in the data.

In summary, we contribute to the current literature on perturbation prediction by demonstrating that GenePert, a combination of a simple prediction model built on top of rich contextual embedding features, outperforms existing models across eight Perturb-seq datasets that are diverse in both the number and the technology used for genetic perturbations. The strength of GenePert lies in its simplicity, scalability, effectiveness, and broad utility.

## 2 Methods

### 2.1 Problem setup and GenePert

Throughout this paper, we will use the following notational conventions. Let *P* denote the total number of *perturbed* genes, and *N*_*p*_ represent the number of cells perturbed with gene *p*; we use *q* to denote the total number of genes for which we measure expression levels. For a given genetic perturbation *p* = 1, …, *P*, we use 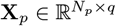 to denote the matrix of expression levels for *q* genes across *N*_*p*_ cells. We adopt the notation *X*_*c*_ and *N*_*c*_ for control cells and **X**_*p*1 +*p*2_ for two-gene perturbations affecting both *p*_1_ and *p*_2_. Finally, we use 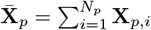 to denote the average gene expression (also referred to as “pseudo-bulk”) of all cells with gene *p* perturbed.

While a few computational methods perturbation effects on a single-cell resolution [Lotfollahi et al., 2020, Dong et al., 2023, Bunne et al., 2023], most prediction and benchmarking efforts have focused on estimating the *average* perturbation effect, defined as 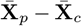. In other words, this can be seen a regression problem where we want to model 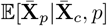 (see Figure 1(1)). The intuition for GenePert is grounded in the biological evidence from prior Perturb-seq experiments that perturbing functionally similar genetic circuits will likely lead to similar post-perturbation expression.

This inspired us to consider the following model for single gene perturbation

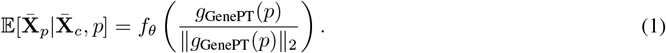

In words, GenePert featurizes each single gene perturbation using the corresponding GenePT gene embedding [Chen and Zou, 2023], denoted as *g*_GenePT_(*p*). GenePT embeddings are large language model embeddings derived from natural language descriptions of gene functions, sourced from NCBI [Schoch et al., 2020] and UniProt [Consortium, 2019], as illustrated in Figure 1(b). The intuition behind using GenePT embeddings is that large language models (LLMs) like GPT-4 have demonstrated remarkable abilities in understanding, reasoning, and even generating biomedical and genomics text [Wei et al., 2024, AI4Science and Quantum, 2023, Hou and Ji, 2024]. As a result, these LLM-derived embeddings of curated gene summaries and functions can directly capture the underlying biology. In Chen and Zou [2023], the authors showed that GenePT embeddings, in particular, excel at biologically relevant prediction tasks, including gene functionality and role prediction, as well as gene-gene and protein-protein interactions across diverse benchmark datasets, highlighting the promise of this approach for Perturb-seq experiments, where prior work often explicitly used gene-gene interaction information [Roohani et al., 2024].

In predicting double gene perturbations, we want GenePert to effectively leverage information from the observed or learned single gene perturbation data and to be permutation-invariant with respect to the inputs of two genes, *p*_1_ and *p*_2_. This motivates the following model:

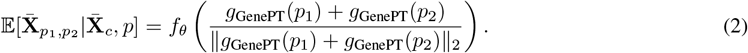

In both (1) and (2), we normalize the input feature to have unit *l*_2_ norm to ensure all features are on the same scale.

Given the relatively high dimensionality of the dataset (with the number of perturbations *P* typically ranging from 20 to 1,000, and an embedding dimension of 3,072 for the largest GenePT embedding), we use ridge regression as *f*_*θ*_ to fit the relationship between the feature and the outcome. Specifically, consider the outcome matrix **X** ∈ ℝ^*P*×*q*^, where the *p*-th row is 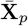, and the feature matrix **G** ∈ ℝ^*P*×*d*^, where the *p*-th row is the 𝓁_2_-normalized *g*_GenePT_(*p*) (or *g*_GenePT_(*p*_1_) + *g*_GenePT_(*p*_2_) for double gene perturbations) and *d* is the embedding dimension. GenePert fits the following ridge regression:

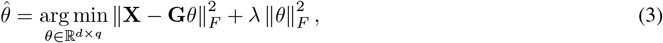

where ∥·∥_*F*_ denotes the Frobenius norm of a matrix, and *λ* > 0 is the tuning parameter. For an unseen genetic perturbation 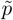 or 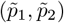 in the test set, the predicted post-perturbation expression is given by 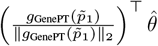 and 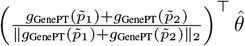, respectively.

### 2.2 Datasets and evaluation metrics

We test our proposed approach using data from eight different perturbation screening datasets across five different cell lines: the K562 cell line [Replogle et al., 2022, Adamson et al., 2016, Norman et al., 2019, Dixit et al., 2016], the retinal pigment epithelial (RPE1) cell line [Replogle et al., 2022], neurons [Tian et al., 2021], the HEK293 cell line [Wessels et al., 2020], and the THP-1 cell line [Xu et al., 2023]. These datasets vary in their tissue of origin and the number of perturbations applied (ranging from 20 to 1,500 unique perturbations). Out of the eight datasets, five were processed and obtained from Roohani et al. [2024] (for the K562 cells and RPE cell line). For the remaining three, we used the following pro-processing procedure to construct the matrix **X** in (3): we first normalize each cell over all genes to have a total sum of 10,000, followed by a log_1+*x*_ transformation. To reduce the complexity of the prediction problem and alleviate the high sparsity in the datasets, we restricted the outcome to only the 5,000 most highly varying genes (that is, *q* = 5, 000), as done in prior work [Roohani et al., 2024, Lotfollahi et al., 2020]. We can compute average expression for each unique perturbation condition to obtain **X**.

In terms of evaluation, we performed five-fold cross-validation on the perturbation space. In each fold, we trained on 80% of the perturbations and evaluated on the remaining 20% of the *unseen* perturbations from the training set. We then aggregated the test set predictions across five folds and reported the prediction performance using both RMSE and Pearson Correlation on the relative scale (i.e., comparing the observed and predicted perturbation effects relative to the average expression of control cells), where PCC stands for the standard Pearson correlation coefficient:

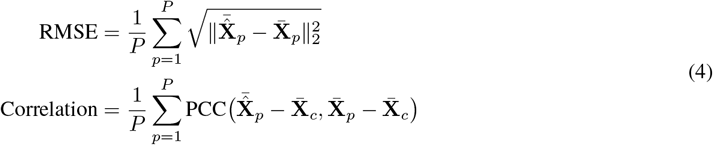

This ensures that all genes have their predictive performance evaluated. We use the GenePT embeddings constructed using GPT-4 embedding models (`text-embedding-3-large` with *d* = 3, 072) in the main GenePert evaluation. We report the best cross-validated metrics across tuning parameters *λ* = 0.1, 1, 10.

We considered the following alternative methods: using the same GenePert setup with ridge regression but varying the embedding vectors, including GenePT embeddings from a smaller embedding model (GPT-3 `text-embedding-ada-002`; *d* = 1, 536), pre-trained Geneformer gene embeddings [Theodoris et al., 2023] (extracted from `Geneformer-6L-30M-i2048`; *d* = 256), pretrained scGPT gene embeddings (extracted from `scGPT` `human`; *d* = 512), and pretrained ESM2 emebddings [Lin et al., 2023] for protein-encoding genes (where we averaged `esm2_t48_15B_UR50D` embeddings of all proteins available for the gene, as done in prior work [Rosen et al., 2024]; *d* = 5, 120). We further tested concatenating ESM2 and GenePT embeddings, as they might offer complementary information on gene sequence and function. We also considered GEARS, a graph neural network-based prediction method that is considered to provide state-of-the-art predictive performance, as well as using the mean expression of the training dataset, which has been demonstrated to perform well on many perturbation datasets in a recent benchmark effort [Kernfeld et al., 2024].

Although GenePert uses ridge regression for *f*_*θ*_ in (1), we explored alternatives like k-nearest neighbors (kNN) with *k* = 1, 5, 10, and a two-layer MLP with hidden dimensions *d* = 256 or 512. However, these options performed less competitively on average compared to ridge regression.

## 3 Results

We report the test set evaluation metrics in (4), aggregated across five-fold cross-validation, in Table 1. We observe that our GenePert approach using GPT-4 gene embeddings consistently outperforms the same approach with other pretrained gene embeddings, achieving the best performance on six out of eight benchmark datasets, measured using RMSE and correlation. The same holds true when comparing GenePert to the simple baseline proposed by Kernfeld et al. [2024], which uses the mean expression of the training set. This result is not surprising, as taking the mean expression of the training set can be viewed as a *special case* of our GenePert approach, where instead of using the gene embeddings *g*_GenePT_(*p*), we include only the intercept term and set the tuning parameter λ = 0.

**Table 1:** Five-fold cross-validated performance on gene expression prediction tasks post-perturbation. We report aggregated test-fold RMSE and correlation (as defined in (4)) across eight Perturb-seq datasets and six different models. The best performances are bolded based on the rounded point estimates.

| Dataset | Embedding | RMSE ($\times 10^{-2}$) ↓ | Correlation ↑ |
| --- | --- | --- | --- |
| Adamson et al. [2016] CRISPRi | GenePert (GPT-4) | **5.30 ± 0.14** | **0.79 ± 0.03** |
| | GenePert (GPT-3.5) | 5.39 ± 0.17 | 0.79 ± 0.04 |
| | ESM2 protein embedding | 5.58 ± 0.23 | 0.78 ± 0.02 |
| | Geneformer | 5.77 ± 0.25 | 0.76 ± 0.03 |
| | scGPT | 5.81 ± 0.29 | 0.76 ± 0.03 |
| | Mean expression of training data | 5.79 ± 0.32 | 0.75 ± 0.03 |
| Dixit et al. [2016] CRISPR-i | GenePert (GPT-4) | 1.85 ± 0.65 | 0.79 ± 0.19 |
|  | GenePert (GPT-3.5) | **1.83 ± 0.40** | **0.80 ± 0.13** |
| | ESM2 protein embedding | 2.04 ± 0.23 | 0.60 ± 0.27 |
| | Geneformer | 2.57 ± 0.80 | 0.29 ± 0.43 |
| | scGPT | 2.84 ± 0.47 | 0.29 ± 0.34 |
| | Mean expression of training data | 3.45 ± 0.20 | 0.19 ± 0.31 |
| Tian et al. [2021] CRISPR-a | GenePert (GPT-4) | **2.54 ± 0.35** | 0.51 ± 0.03 |
| | GenePert (GPT-3.5) | **2.54 ± 0.34** | 0.51 ± 0.03 |
| | ESM2 protein embedding | 2.56 ± 0.34 | 0.51 ± 0.03 |
| | Geneformer | 2.56 ± 0.34 | 0.51 ± 0.03 |
| | scGPT | 2.56 ± 0.34 | 0.51 ± 0.03 |
| | Mean expression of training data | 2.56 ± 0.34 | 0.51 ± 0.03 |
| Replogle et al. [2022] (K562, CRISPR-i) | GenePert (GPT-4) | **7.02 ± 0.14** | **0.51 ± 0.01** |
| | GenePert (GPT-3.5) | 7.17 ± 0.15 | 0.49 ± 0.01 |
| | ESM2 protein embedding | 7.40 ± 0.13 | 0.46 ± 0.01 |
| | Geneformer | 7.68 ± 0.12 | 0.42 ± 0.01 |
| | scGPT | 7.51 ± 0.12 | 0.44 ± 0.01 |
| | Mean expression of training data | 7.90 ± 0.11 | 0.38 ± 0.01 |
| Norman et al. [2019] CRISPR-i | GenePert (GPT-4) | **3.56 ± 0.23** | **0.79 ± 0.04** |
| | GenePert (GPT-3.5) | 4.05 ± 0.21 | 0.75 ± 0.03 |
| | ESM2 protein embedding | 3.90 ± 0.19 | 0.75 ± 0.04 |
| | Geneformer | 4.54 ± 0.20 | 0.69 ± 0.03 |
| | scGPT | **3.56 ± 0.17** | 0.76 ± 0.03 |
| | Mean expression of training data | 5.78 ± 0.28 | 0.50 ± 0.02 |
| Replogle et al. [2022] (RPE1, CRISPR-i) | GenePert (GPT-4) | **9.57 ± 0.07** | **0.65 ± 0.01** |
| | GenePert (GPT-3.5) | 9.67 ± 0.09 | 0.64 ± 0.00 |
| | ESM2 protein embedding | 9.85 ± 0.10 | 0.63 ± 0.00 |
| | Geneformer | 10.10 ± 0.18 | 0.63 ± 0.00 |
| | scGPT | 9.96 ± 0.17 | 0.63 ± 0.00 |
| | Mean expression of training data | 10.36 ± 0.17 | 0.63 ± 0.00 |
| Wessels et al. [2020] (CRISPR-Cas13d) | GenePert (GPT-4) | **3.71 ± 0.12** | **0.76 ± 0.03** |
| | GenePert (GPT-3.5) | 3.72 ± 0.12 | **0.76 ± 0.03** |
| | ESM2 protein embedding | 3.84 ± 0.13 | 0.75 ± 0.03 |
| | Geneformer | 3.91 ± 0.14 | 0.73 ± 0.03 |
| | scGPT | 3.78 ± 0.14 | 0.75 ± 0.03 |
| | Mean expression of training data | 4.81 ± 0.14 | 0.62 ± 0.03 |
| Xu et al. [2023] (CRISPR-dCas9) | GenePert (GPT-4) | **2.37 ± 0.21** | **0.34 ± 0.02** |
| | GenePert (GPT-3.5) | 2.38 ± 0.21 | 0.33 ± 0.02 |
| | ESM2 protein embedding | 2.38 ± 0.22 | 0.32 ± 0.02 |
| | Geneformer | 2.39 ± 0.21 | 0.31 ± 0.02 |
| | scGPT | 2.39 ± 0.21 | 0.31 ± 0.02 |
| | Mean expression of training data | 2.37 ± 0.22 | 0.33 ± 0.02 |

Additionally, GenePert also outperformed GEARS by a meaningful margin on datasets with a large number of perturbations: GEARS reported 15% held-out test set correlations of 0.52, 0.28, and 0.45 on the Replogle RPE1, Replogle K562, and Norman datasets, respectively, all of which are meaningfully lower than the metrics reported by our approach in Table 1. On three datasets (Adamson, Replogle RPE1, and Replogle K562), we also compared our performance with a linear model using principal components of observed gene expressions, as implemented in Ahlmann-Eltze et al. [2024]. The linear model’s performance, when aggregated across folds, is similar to the mean expression baseline of the training data but lags behind GenePert.

We also evaluated the effectiveness of the simple additive assumption in (2) by assessing the predictive performance stratified by combinations of gene perturbations in Table 3. Similar to our observations in the single or pooled perturbation cases, the GenePert approach often achieves the best results.

Encouraged by the performance of GenePert, we further investigated whether ensembling embeddings derived from different sources (e.g., sequence-based models like ESM2 or scGPT and GenePT) could further improve the predictive performance for unseen perturbations. The results of concatenating GenePT and ESM2 embeddings are presented in Table 2, suggesting that ensembling in this context provides a small benefit when evaluated using correlation.

**Table 2:** Combining GenePT and ESM2 embeddings increases the five-fold cross-validated prediction metrics marginally. We investigated whether (i) concatenating GenePT and ESM2 embeddings and (ii) using GPT embeddings of gene names only could improve test set RMSE and correlation (as defined in (4)) across Perturb-seq datasets. Overall, all methods perform similarly, with a small improvement observed when evaluated using correlation when ensembling different embeddings. Full GenePT embeddings outperform the ablated version with gene names only on most datasets, but the performances are broadly similar.

| Dataset | Embedding | RMSE ($\times 10^{-2}$) ↓ | Correlation ↑ |
| --- | --- | --- | --- |
| Adamson et al. [2016] CRISPRi | GenePert (GPT-4) | $5.30 \pm 0.14$ | $0.79 \pm 0.03$ |
| | Gene Name Embedding (GPT-4) | $5.42 \pm 0.16$ | $0.78 \pm 0.04$ |
|  | Concatenated GenePT (GPT-4) and ESM2 Embedding | **$5.29 \pm 0.10$** | **$0.81 \pm 0.03$** |
| Dixit et al. [2016] CRISPR-i | GenePert (GPT-4) | $1.85 \pm 0.65$ | **$0.79 \pm 0.19$** |
| | Gene Name Embedding (GPT-4) | $1.80 \pm 0.66$ | $0.66 \pm 0.16$ |
| | Concatenated GenePT (GPT-4) and ESM2 Embedding | **$1.58 \pm 0.57$** | $0.76 \pm 0.11$ |
| Tian et al. [2021] CRISPR-a | GenePert (GPT-4) | **$2.54 \pm 0.35$** | **$0.51 \pm 0.03$** |
| | Gene Name Embedding (GPT-4) | $2.55 \pm 0.34$ | **$0.51 \pm 0.03$** |
|  | Concatenated GenePT (GPT-4) and ESM2 Embedding | **$2.54 \pm 0.35$** | **$0.51 \pm 0.03$** |
| Replogle et al. [2022] (K562, CRISPR-i) | GenePert (GPT-4) | $7.02 \pm 0.14$ | $0.51 \pm 0.01$ |
| | Gene Name Embedding (GPT-4) | $7.28 \pm 0.15$ | $0.46 \pm 0.01$ |
|  | Concatenated GenePT (GPT-4) and ESM2 Embedding | **$6.99 \pm 0.13$** | **$0.52 \pm 0.01$** |
| Norman et al. [2019] CRISPR-i | GenePert (GPT-4) | $3.56 \pm 0.23$ | $0.79 \pm 0.04$ |
|  | Gene Name Embedding (GPT-4) | **$3.52 \pm 0.13$** | **$0.88 \pm 0.01$** |
| | Concatenated GenePT (GPT-4) and ESM2 Embedding | $3.60 \pm 0.24$ | $0.80 \pm 0.04$ |
| Replogle et al. [2022] (RPE1, CRISPR-i) | GenePert (GPT-4) | $9.57 \pm 0.07$ | **$0.65 \pm 0.01$** |
| | Gene Name Embedding (GPT-4) | $9.87 \pm 0.12$ | $0.64 \pm 0.00$ |
|  | Concatenated GenePT (GPT-4) and ESM2 Embedding | **$9.50 \pm 0.06$** | **$0.65 \pm 0.01$** |
| Wessels et al. [2020] (CRISPR-Cas13d) | GenePert (GPT-4) | **$3.71 \pm 0.12$** | **$0.76 \pm 0.03$** |
| | Gene Name Embedding (GPT-4) | $3.71 \pm 0.12$ | $0.75 \pm 0.03$ |
| | Concatenated GenePT (GPT-4) and ESM2 Embedding | $3.72 \pm 0.13$ | **$0.76 \pm 0.03$** |
| Xu et al. [2023] (CRISPR-dCas9) | GenePert (GPT-4) | **$2.37 \pm 0.21$** | **$0.34 \pm 0.02$** |
| | Gene Name Embedding (GPT-4) | $2.37 \pm 0.22$ | $0.33 \pm 0.02$ |
| | Concatenated GenePT (GPT-4) and ESM2 Embedding | **$2.37 \pm 0.21$** | $0.33 \pm 0.02$ |

**Table 3:** GenePert provides best results for gene expression prediction in double gene perturbations. We report aggregated test-fold RMSE and correlation (as defined in (4)) on the subset of double gene perturbations from two Perturb-seq datasets. We note that GenePert (or combining GenePert and ESM2 embeddings) provides the best RMSE and correlation on both datasets.

| Dataset | Embedding | RMSE ($\times 10^{-2}$) ↓ | Correlation ↑ |
| --- | --- | --- | --- |
| Norman et al. [2019] CRISPR-i | GenePert (GPT-4) | $3.50 \pm 0.14$ | **$0.85 \pm 0.02$** |
| | GenePert (GPT-3.5) | $4.22 \pm 0.16$ | $0.80 \pm 0.02$ |
| | ESM2 protein embedding | $3.83 \pm 0.15$ | $0.77 \pm 0.03$ |
| | Geneformer | $4.63 \pm 0.20$ | $0.71 \pm 0.02$ |
| | scGPT | **$3.46 \pm 0.15$** | $0.79 \pm 0.03$ |
| | Mean expression of training data | $6.16 \pm 0.23$ | $0.53 \pm 0.02$ |
| | Concatenated GenePT (GPT-4) and ESM2 Embedding | $3.55 \pm 0.14$ | **$0.85 \pm 0.02$** |
| Wessels et al. [2020] CRISPR-Cas13d | GenePert (GPT-4) | **$3.53 \pm 0.10$** | $0.78 \pm 0.01$ |
| | GenePert (GPT-3.5) | $3.53 \pm 0.11$ | $0.80 \pm 0.02$ |
| | ESM2 protein embedding | $3.63 \pm 0.12$ | $0.78 \pm 0.02$ |
| | Geneformer | $3.72 \pm 0.15$ | $0.77 \pm 0.02$ |
| | scGPT | $3.65 \pm 0.12$ | $0.78 \pm 0.02$ |
| | Mean expression of training data | $4.76 \pm 0.07$ | $0.65 \pm 0.01$ |
| | Concatenated GenePT (GPT-4) and ESM2 Embedding | $3.54 \pm 0.10$ | **$0.80 \pm 0.01$** |

Furthermore, to investigate the proportion of training perturbations needed to achieve good test set performance, we plot the test set metric performance as a function of the proportion of unique gene perturbations used for training in Figure 2. We note that (i) ridge regression models outperform KNN regression models on average, and (ii) ridge regressions achieve high test set correlations even with a low number of perturbations used for training. For instance, the regression model with optimal tuning parameters outperformed baseline methods at 10-20% (around 100-200 perturbations in the RPE1 and K562 datasets). This further highlights the sample efficiency brought by the GenePert approach by leveraging the biological knowledge encoded in the GenePT embeddings.

**Figure 2:**
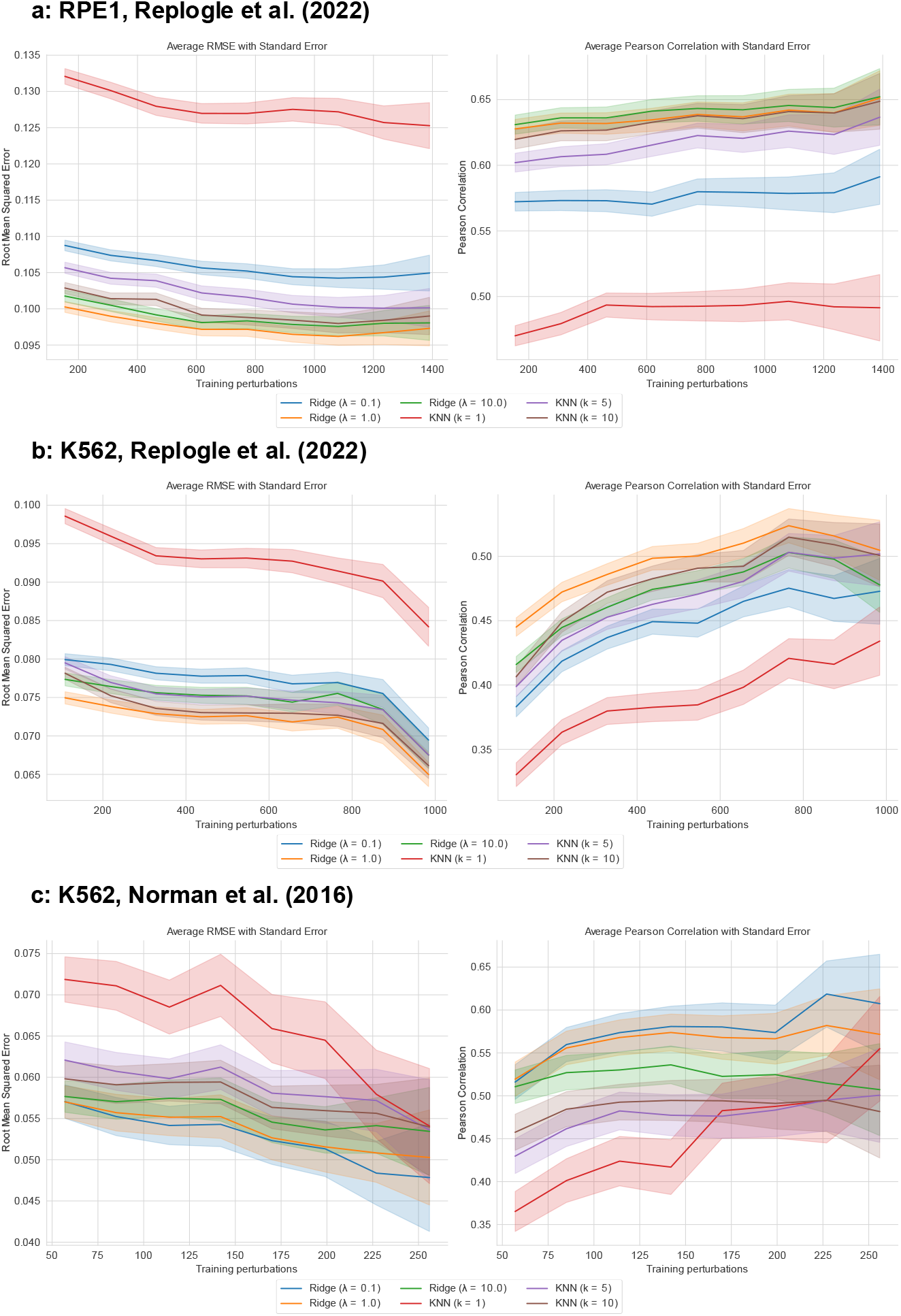
GenePert is sample-efficient in generating state-of-the-art predictions across datasets. We display GenePert results across six simple prediction models *f*_*θ*_ (each represented by a line: three ridge regression models and three kNN models with different tuning parameters) on (a) the RPE1 dataset from Replogle et al. [2022], (b) the K562 dataset from Replogle et al. [2022], and (c) the K562 dataset from Norman et al. [2019]. In each panel, we plot test set performance (left: RMSE, right: Pearson correlation as defined in (4)) as a function of the proportion of unique perturbations used for training. We observe that (i) across all models, GenePert often achieves near-optimal performance with around 20% of the perturbations used for training, validating its sample efficiency, and (ii) ridge regression generally outperforms kNN regression.

To understand the competitive performance of GenePert, we further visualize the test set metrics for individual perturbations in Figure 3 for the K562 datasets collected by Replogle et al. [2022]. Panel (a) displays the test set correlation as a function of the smallest cosine distance between the test perturbation and training perturbations in the GenePT embedding space. We observe that as the distance increases, the test set correlation tends to decrease, aligning with the intuition that more dissimilar genetic perturbations are harder to extrapolate. Additionally, in Figure 3(b) and (c), we observe that while the GenePert approach achieves noticeably better performance in aggregate, there are subsets of genes where alternative methods perform well, suggesting a promising ensembling approach. We have also tested the results of ablating the gene summary information and using GPT-4 text embeddings for *gene names only* as input. This approach achieves competitive results (see Table 2) but, as expected, lags slightly behind the GenePT embeddings that use the full context, suggesting the robustness of this approach.

**Figure 3:**
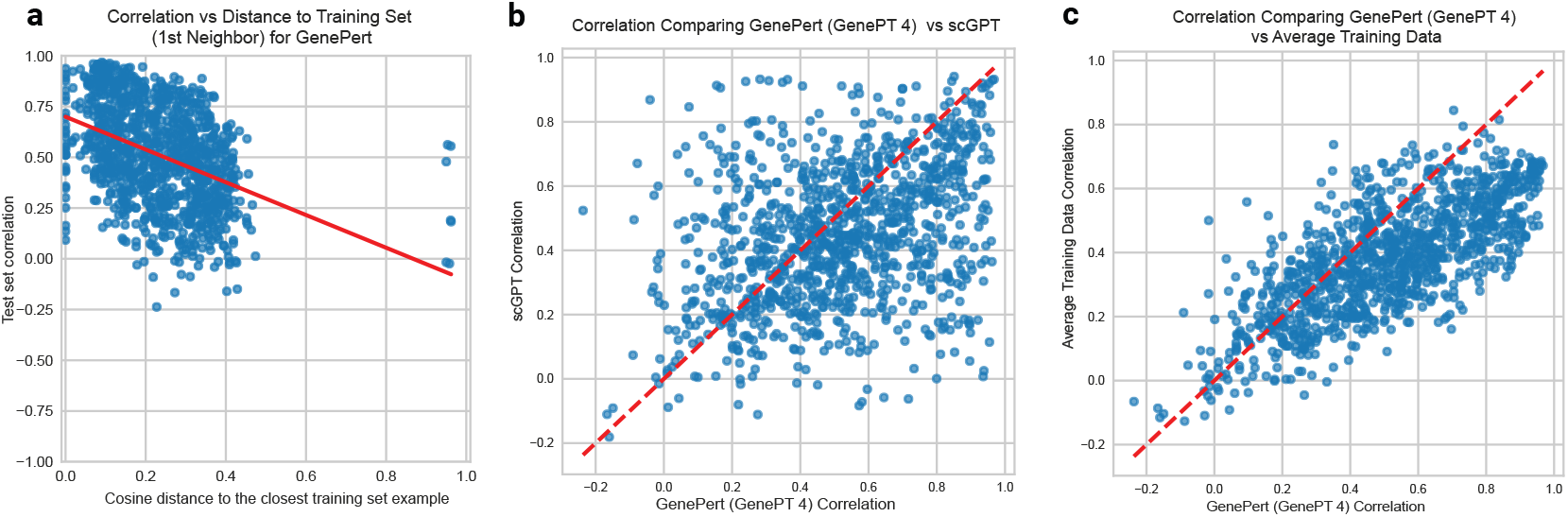
Gene expression prediction performance on the individual gene perturbation level on the K562 dataset collected by Replogle et al. [2022]. (a): There is a negative correlation (Pearson *r* = 0.41, *p* < 0.001) between the distance to the closest training example and the test set correlation of the gene. (b): Scatter plot of the test correlation of the GenePert (GPT-4) versus using scGPT embeddings; dashed line indicates *y* = *x*. (c): same as (b), but for GenePert (GPT-4) versus training set average.

## 4 Discussion

With the advancement of technologies to perform genetic and cellular perturbation studies at scale, there is increasing interest in deciphering the underlying genetic causes and molecular mechanisms from these studies. In particular, recent advances in computational models, such as large-scale deep learning models [Theodoris et al., 2023, Cui et al., 2024, Hao et al., 2024, Roohani et al., 2024], have shown promise in predicting perturbations and their combinations at scale, especially for unseen perturbations. However, training and fine-tuning these models can be sample-inefficient, especially when the number of unique perturbations is limited in available datasets.

In this work, we propose GenePert, a simple ridge regression model based on GenePT embeddings, which achieves state-of-the-art prediction accuracy on benchmark datasets widely used for Perturb-seq across cell lines and perturbation mechanisms. Additionally, GenePert is both sample-efficient (achieving state-of-the-art correlation metrics with less than 20% of perturbations used for training) and easy to extend to unseen genes (requiring only the GenePT embedding of the unseen gene perturbation). Combining sequence-based embeddings such as ESM2 embeddings with text-based GenePT embeddings also provides improvements in several datasets, suggesting that these embeddings offer complementary information for downstream biological tasks, as independently observed by Märtens et al. [2024].

It is important to note the limitations of GenePert, as it relies solely on available gene summaries and descriptions, which may overlook the unknown regulatory roles of perturbed genes. Furthermore, the embeddings used in GenePert might not fully capture the context-dependent roles of genes and cells in specific tissues and cell types, as they are derived from pretrained large language models. While these embeddings demonstrate empirical effectiveness, exploring opportunities to fine-tune the underlying language model directly could improve prediction accuracy for more complex perturbations and unseen cell types or contexts. Moreover, while the average gene expression for each perturbation is of primary interest in perturbation screens, our approach only estimates a common perturbation effect and does not account for individual cell effects.

Several promising directions lie ahead for future research. First, extending the current GenePert approach to be more dynamic and context-dependent could enhance its utility in real-world applications. For instance, this would involve understanding the transferability of trained models from one dataset to another, within or even across cell lines [Ji et al., 2024, Mourragui et al., 2021]. Additionally, it is natural to assess whether GenePert could be generalized to other modalities of perturbations, such as drugs and small molecules [Lotfollahi et al., 2023, 2019, Srivatsan et al., 2020], as well as generating predictions beyond RNA expression outcomes to protein levels [Papalexi et al., 2021, Frangieh et al., 2021] and disease phenotype predictions [Yao et al., 2023] under perturbation.

